# Mechanistic insights into RNA cleavage by bacterial RNA polymerase from a comprehensive mutational screen

**DOI:** 10.1101/2024.06.20.599782

**Authors:** Janne J. Mäkinen, Georgiy A. Belogurov

**Affiliations:** University of Turku, Department of Life Technologies, Turku, Finland

**Keywords:** RNA polymerase/proofreading/backtracking/RNA cleavage

## Abstract

RNA polymerase (RNAP) mediates the synthesis of an RNA copy of the template DNA—the first and often decisive step in gene expression. All cellular RNAPs possess an additional capacity to cleave nucleotides from the 3’ end of the nascent RNA. This ability potentially enhances the efficiency and accuracy of transcription, enabling RNAP to maintain processivity and ensure the fidelity of the RNA transcript. This study investigates the contributions of various active site regions to the RNA cleavage activity using a large collection of *Escherichia coli* RNAP variants. Unlike previous studies conducted under non-physiological conditions, this research employed backtracked RNAP complexes that cleave nascent RNA on a timescale of minutes under physiological pH and low Mg^2+^ concentrations. Our findings provide key insights into the RNA cleavage activity of the RNAP active site. Complete closure of the active site by the Trigger Loop (TL) facilitates RNA cleavage in 1-nt backtracked states, but not in 2-nt backtracked states. However, the RNA-proximal N-terminus of the TL influences the cleavage rate in both states. β subunit Asp814 plays an important role in RNA cleavage, regardless of backtracking depth, likely by coordinating the Mg^2+^ ion responsible for generating the nucleophile. During RNA cleavage, the pre-translocated RNA nucleotide is base-paired to the template DNA, but its sugar-phosphate backbone is shifted compared to canonical pre-translocated and NTP-bound states. Bulky substitutions in the E-site (NTP entry area) stimulate RNA cleavage, suggesting that RNA binding in this site inhibits the reaction.

## Introduction

RNA polymerase (RNAP) mediates the synthesis of an RNA copy of the template DNA—the first and often decisive step in gene expression. All RNAPs transcribing cellular genomes are multisubunit enzymes with conserved catalytic cores. The bacterial RNAP, comprising five subunits, ααββ’ω, is the simplest model system used for studying the fundamental mechanistic properties of all multisubunit RNAPs. During transcription, RNAP separates approximately 10-12 base pairs of the DNA template, while the nascent RNA forms a 9-11 bp hybrid with the template DNA. This structure, called the transcription bubble, is unstable without RNAP, necessitating processive synthesis of the entire transcript without dissociation from the DNA template. To ensure processivity and transcription accuracy, RNAPs have evolved a proofreading capability. These enzymes can detect mismatches between the 3’ end of the nascent RNA and the DNA template. In such cases, RNAP backtracks along the DNA template, cleaves erroneous RNA nucleotides from the 3’ end, and resumes transcription from the newly generated 3’ end of the nascent RNA.

RNAPs maintain high fidelity by selecting the correct nucleobase with a preference ratio ranging from several hundred to several thousand-fold (Yuzenkova *et al*, 2010; Chung *et al*, 2023; Gout *et al*, 2013; Imashimizu *et al*, 2015). Upon misincorporation the mismatched nucleotide frays and significantly slows the addition of the next nucleotide, thereby pausing transcription (Sydow *et al*, 2009). When RNA elongation pauses, RNAP can backtrack along the DNA (Komissarova & Kashlev, 1997b, 1997a; Nudler *et al*, 1997), extruding the nascent RNA into the substrate loading channel (secondary channel) (Wang *et al*, 2009; Cheung & Cramer, 2011; Abdelkareem *et al*, 2019; Sekine *et al*, 2015; Wee *et al*, 2023). Backtracking is not solely dependent on misincorporation; any slowdown of RNA synthesis or impediment to forward translocation, such as a roadblock, can cause RNAP to backtrack. The propensity for backtracking is influenced by the transcribed sequence, and mismatches at the 3’ end of the RNA greatly facilitate backtracking.

This extruded RNA can be cleaved either by the intrinsic endonuclease activity of the RNAP active site (Orlova *et al*, 1995; Zenkin *et al*, 2006; Esyunina *et al*, 2016; Sydow *et al*, 2009; Kireeva *et al*, 2008; Mishanina *et al*, 2017) or more efficiently with auxiliary factors (Erie *et al*, 1993; Sydow & Cramer, 2009; Reines, 1992; Bubunenko *et al*, 2017; Borukhov *et al*, 1993; Laptenko *et al*, 2003; Hausner *et al*, 2000; Izban & Luse, 1992) and specialized subunits (Ruan *et al*, 2011). Multi-subunit RNAPs use the same catalytic site for both RNA synthesis and RNA hydrolysis. The accessory cleavage factors and subunits do not feature a dedicated active site on their own but merely contribute several amino acid residues to the RNAP active site thereby switching it from synthetic to hydrolytic activity. The RNA is always cleaved between the pre- and post-translocation register, restoring the catalytically proficient post-translocated configuration of the RNAP active site. RNAP nearly always cleaves an RNA fragment that is one nucleotide longer than the mismatched segment because the cleavage occurs efficiently only between the two nucleotides that base-pair with the template DNA. RNA cleavage likely involves two Mg^2+^ ions (Sosunova *et al*, 2013). A tightly bound Mg^2+^ ion #1 is coordinated by universally conserved aspartate triad (Asp 460, 462 and 464 in *Escherichia coli* RNAP) and participates in both RNA cleavage and RNA synthesis reactions. The weakly bound and never directly observed Mg^2+^ ion #2 is likely located differently from the Mg^2+^ ion #2 brought in by the substrate NTP during RNA synthesis reaction (Sosunova *et al*, 2013).

RNA cleavage has been previously studied by many research teams. Some studies have explored the role of catalytic Mg^2+^ ions and the mechanistic relationships between nucleotide addition and RNA cleavage reactions (Sosunov *et al*, 2003; Zenkin *et al*, 2006; Sosunova *et al*, 2013). Others have focused on the mobile element of the active site, known as the Trigger Loop (TL) and its involvement in RNA cleavage and backtracking (Yuzenkova & Zenkin, 2010; Zhang *et al*, 2010; Esyunina *et al*, 2016; Mishanina *et al*, 2017; Turtola *et al*, 2018; Mosaei & Zenkin, 2021).

Additionally, several studies have measured the RNA cleavage activities of *E. coli* RNAPs with amino acid substitutions mimicking more cleavage-proficient RNAPs from other bacterial species (Esyunina *et al*, 2016; Riaz-Bradley *et al*, 2020). These diverse approaches have contributed to our understanding of RNA cleavage and the mechanisms underlying this process.

In this study we employed a collection of 28 *E. coli* RNAP variants to elucidate the contributions of different regions of the active site to the RNA cleavage reaction. Unlike previous studies conducted at high pH and Mg^2+^ concentration, this work utilized backtracked complexes that cleave the nascent RNA on a timescale of minutes under physiological pH and low Mg^2+^ concentration. Our results delineate the optimal geometry of backtracked RNA for the cleavage reaction and elucidate the role of the TL and the E-site in RNA cleavage at varying depths of backtracking.

## Results and Discussion

The rate of RNA cleavage is known to be affected by the transcribed sequence. We chose two nucleic acid scaffolds from a library of over 200 available in our laboratory to assemble backtracked complexes that can be nearly completely cleaved by the wild-type *E. coli* RNAP in one hour at 25 ℃, pH 7.5, and 2 mM Mg^2+^ (**Figure 1**). This selection enabled us to carry out RNA cleavage assays within a reasonable timeframe while maintaining the stability of RNAP in the reaction mixture. The 1-nt backtracked complex (1BKT) was predicted to adopt a state where the penultimate RNA nucleotide occupied the pre-translocated register (substrate site, A-site) and the 3’ terminal RNA nucleotide protruded into the secondary channel, due to a single mismatched base at the RNA 3’ end. Similarly, the 2-nt backtracked complex (2BKT) was predicted to adopt a state where the third nucleotide from the RNA 3’ termini occupied the pre-translocated register and the two 3’ terminal RNA nucleotides were extruded into the secondary channel, because of two mismatched RNA bases at the RNA 3’ end.

**Figure 1.**
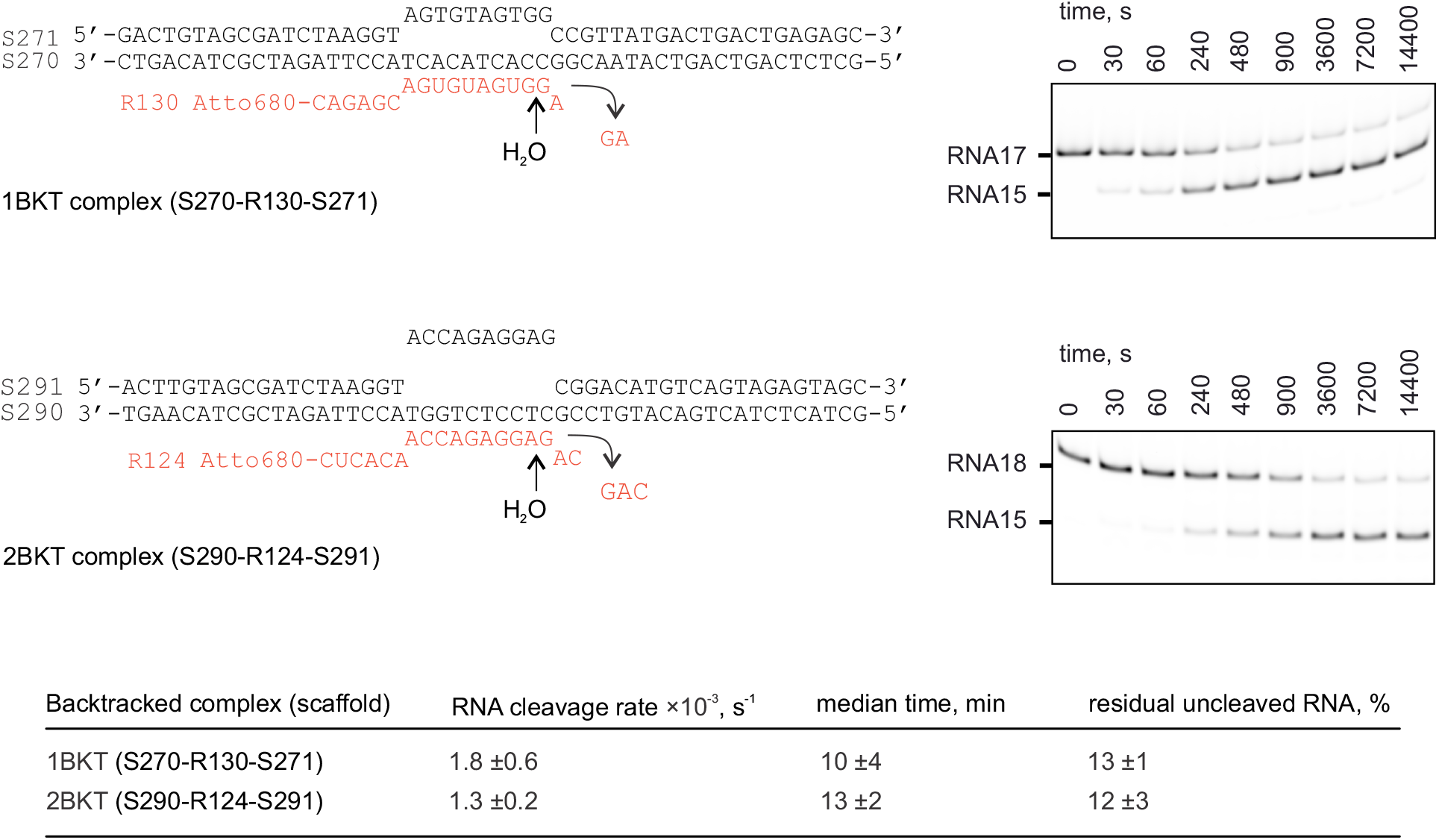
RNA cleavage in 1BKT and 2BKT by the wild-type *E. coli* RNAP.

We then measured the rate of RNA cleavage in 1BKT and 2BKT using a large collection of *E. coli* RNAP variants available in the laboratory. The type of amino acid substitutions that we used in our experiments varied greatly. Sometimes, both alanine and more conservative substitutions of active site residues were available and tested (e.g., βE813A and βE813Q). In other cases, the mutations mirrored variations in the active site structure between RNAPs from different species (e.g., β’Q504R, β’Y457F). Another subset of substitutions located at the periphery of the active site was chosen because they stimulate the closure of the RNAP active site (β’F773V, β’P750L, β’G1136S) (Malinen *et al*, 2014; Turtola *et al*, 2018). Finally, some substitutions, such as β’K598W, had complex design rationales but strongly affected RNA cleavage activity in our previous studies (Turtola *et al*, 2018). **Figures 2-3** summarize the effects of amino acid residue substitutions on the RNA cleavage rate.

**Figure 2.**
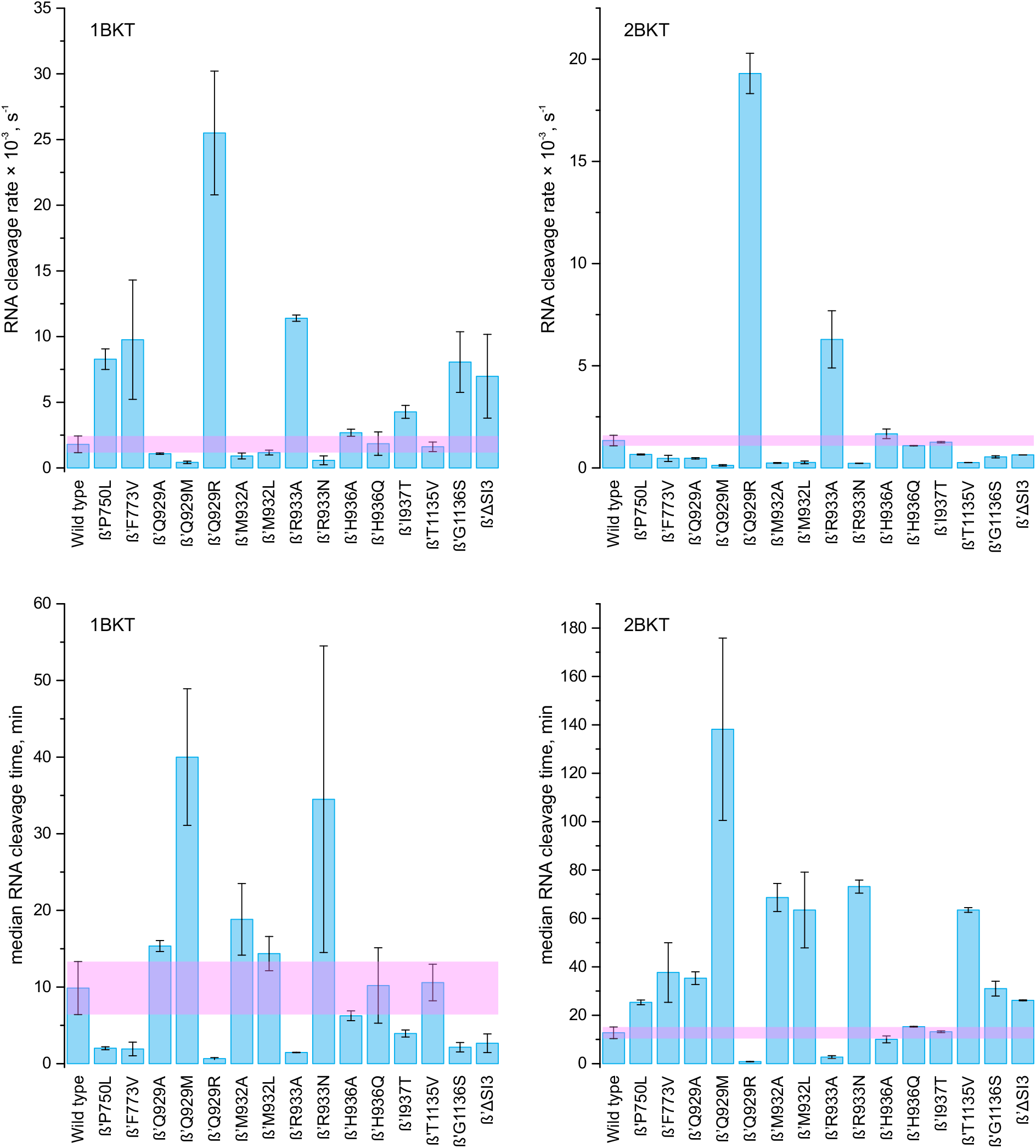
RNA cleavage activities of *E. coli* RNAPs with amino acid substitutions in TL and adjacent regions that affect folding of TL into TH. The horizontal magenta bar depicts the RNA cleavage activity of the wild-type RNAP ±SD. Top graphs accentuate fast cleaving enzymes, whereas bottom graphs accentuate slow cleaving enzymes.

**Figure 3.**
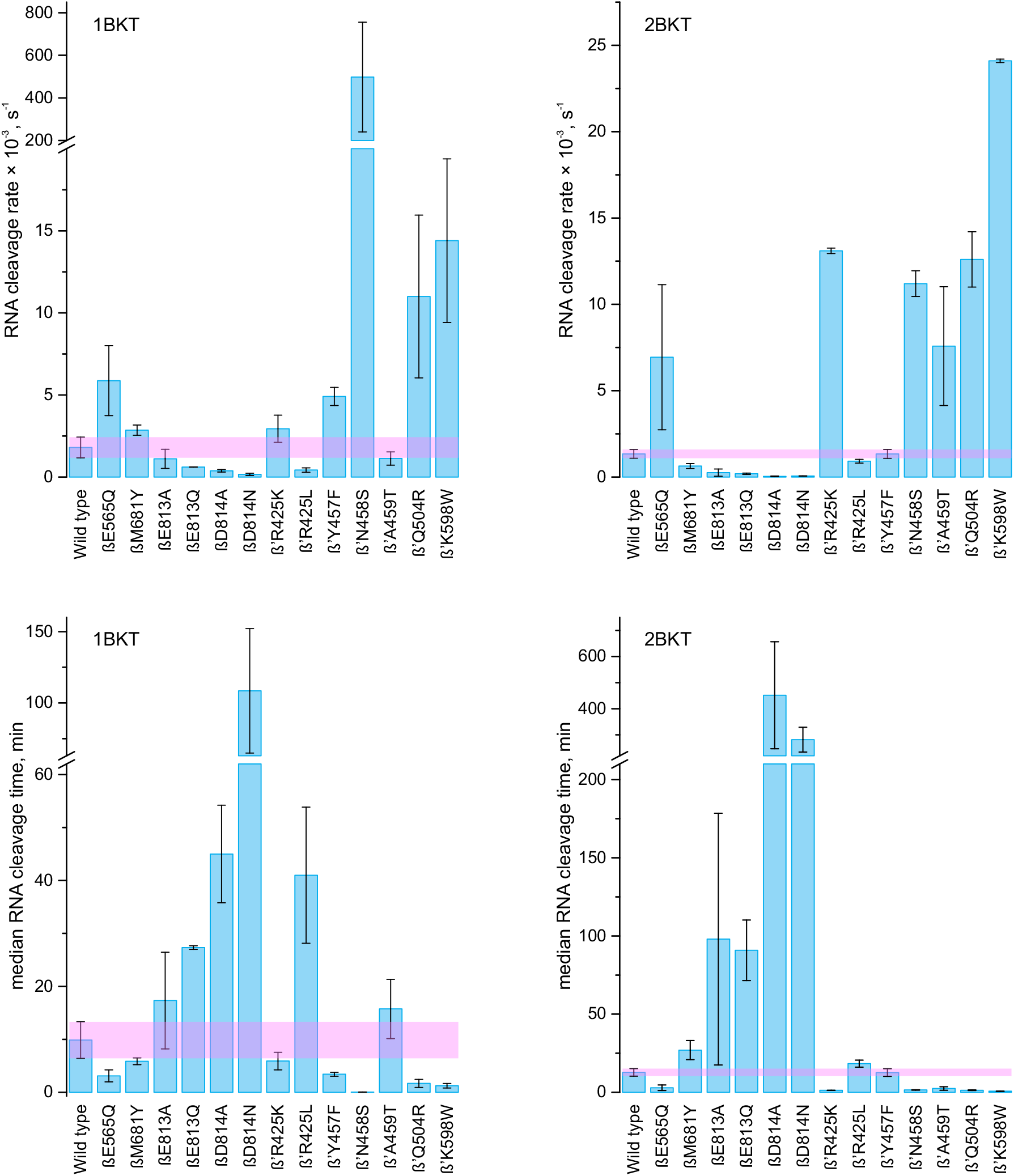
RNA cleavage activities of *E. coli* RNAPs with amino acid substitutions in the main active site cavity and the E-site. The horizontal magenta bar depicts the RNA cleavage activity of the wild-type RNAP ±SD. Top graphs accentuate fast cleaving enzymes, whereas bottom graphs accentuate slow cleaving enzymes.

### The role of the TL domain in catalysis of the nascent RNA cleavage

During the catalysis of the nucleotide addition by the multisubunit RNAPs, the substrate NTP initially binds to the active site in the inactive conformation. This binding is followed by the closure of the active site by a mobile element called the TL, aligning the NTP for catalysis (Vassylyev *et al*, 2007). In gram-positive bacteria, eukaryotes, and archaea, the TL is relatively small and alternates between open (loop-like conformations) and closed (two-helical bundle conformation or Trigger Helices; TH) states. The TH conformation is stabilized by interactions with the Bridge Helix (BH), a long helical element that bridges the cleft between β and β’ subunits, and the F-loop, an N-terminal extension of the BH. Notably, in the majority of gram-negative bacteria, the TL contains a large insertion known as SI3 (**Figure 4A**). This addition affects the dynamics of TL to TH transition, with SI3 inhibiting TL folding into TH in certain sequence contexts (Bao & Landick, 2021; Bao *et al*, 2024), but having little effect in others (Agapov *et al*, 2020).

**Figure 4.**
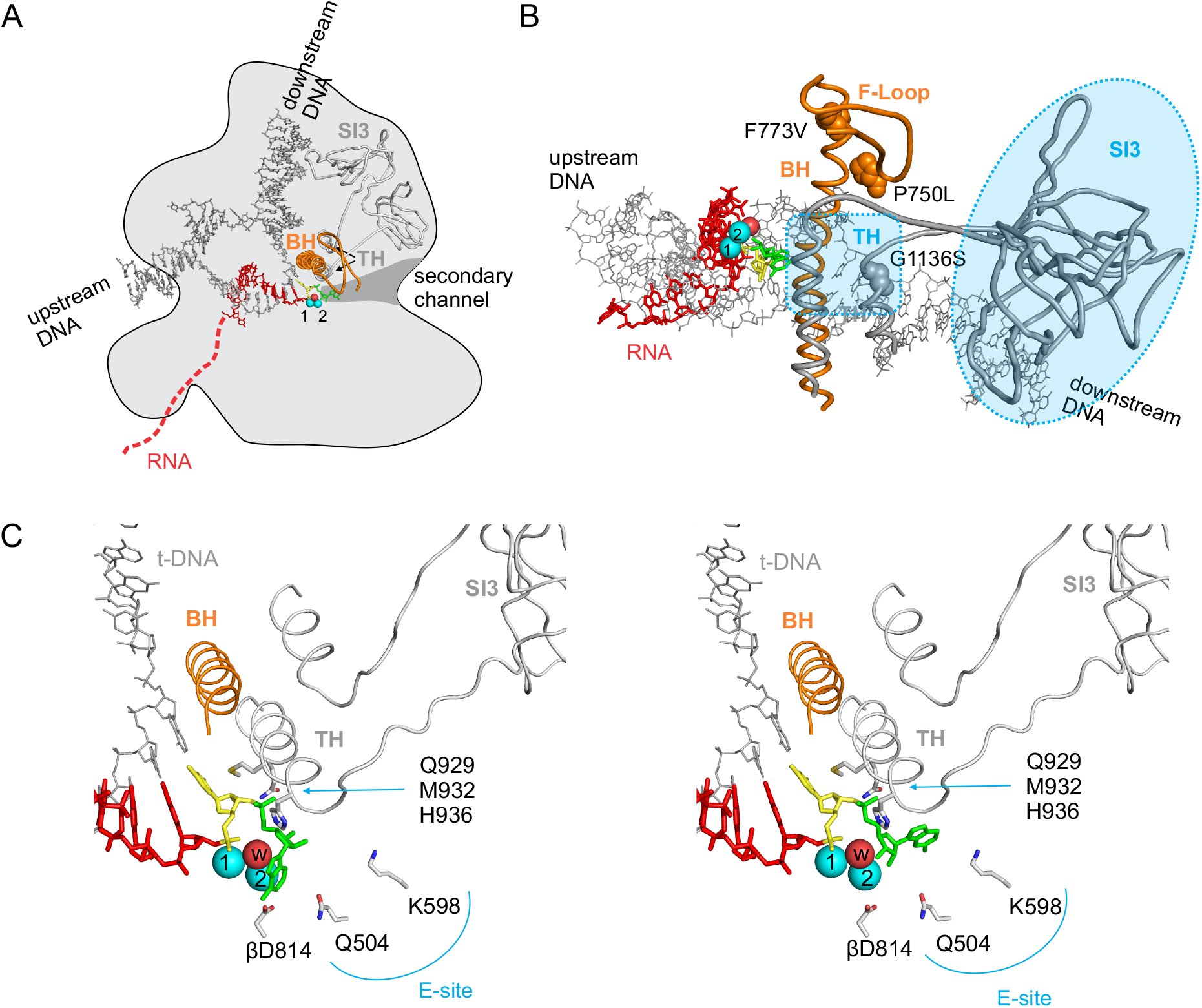
Structural models of 1-nt backtracked state of bacterial RNAP. 1-nt backtracked state likely permits the complete folding of the TL into TH (2 helical turns) **(A)** A top view depicts the transcription bubble, the secondary channel that delivers substrate NTP and accommodate the backtracked nucleotides, the prominent elements of the active site called the Bridge Helix (BH, orange) and the TH (gray). The N-terminal extension of the BH called F-loop is also shown in orange. The large insertion domain in the TL called SI3 that is present in most gram-negative bacteria and absent in gram-positive bacteria, Eukaryotes and Archaea is shown in gray. Backtracked RNA nucleotide is colored green, nucleotide residing in the pre-translocated register is colored yellow. Mg^2+^ ions are shown as cyan spheres. Mg^2+^ ion #1 is tightly bound and crystallographically observed in most RNAP structures, Mg^2+^ #2 is weakly bound, is not crystallographically observed, and is modeled to arrive at a plausible geometry for RNA cleavage reaction. A nucleophilic water molecule coordinated by Mg^2+^ #2 is also modeled in and depicted as a red sphere. **(B)** A side view depicting amino acid substitutions known to stabilize the folded helical state of the TL. **(C)** The backtracked nucleotide has been proposed to fold back towards the Mg^2+^ ions in the cleavage-proficient conformation (left, Sosunova et al 2013) or occupy the E-site (a loosely defined area near β’Gln504 and β’Lys598) in the cleavage-deficient conformation (right, Turtola et al, 2018).

The effect of TL on RNA cleavage is a subject of debate. In certain systems, TL significantly stimulates the cleavage reaction (Yuzenkova & Zenkin, 2010; Esyunina *et al*, 2016), whereas in others, it has a negligible effect (Zhang *et al*, 2010). Biochemical studies involving structural modeling suggest that the closed RNAP active site can accommodate one backtracked nucleotide (Turtola *et al*, 2018). TH folding can stabilize the complex in a 1-nt backtracked state and facilitate RNA cleavage by positioning the backtracked nucleotide in the active site (Mishanina *et al*, 2017; Turtola *et al*, 2018). However, two backtracked nucleotides cannot fit into the closed active site, suggesting that TH should not improve the stability of the 2-nt backtracked state. However, TL may still contribute to catalysis of RNA cleavage in an unfolded or partially folded conformation.

Our findings demonstrate that substitutions favoring TH folding (β’F773V, β’P750L, β’G1136S, ΔSI3) enhanced RNA cleavage in 1BKT (∼5-fold) (**Figure 2, 4B**) but had a negative effect (2-3-fold) on cleavage in 2BKT (**Figure 2, 5**). This suggests that TH folding promotes cleavage in 1BKT by stabilizing the 1-nt backtracked state and positioning the RNA for efficient cleavage. Interestingly, substitutions located in the second helical turn of the folded TH (β’H936A, β’H936Q, β’I937T) (**Figure 4C**) had little effect on RNA cleavage in our system, despite they were reported to have large effects in other systems (Yuzenkova & Zenkin, 2010). At the same time, substitutions of β’Gln929 and β’Arg933 in the first helical turn of TH had relatively large effects on cleavage in 1BKT. The complete folding of TH is incompatible with the 2-nt backtracked state due to clashes with the backtracked RNA (**Figure 5**). As a result, the complete folding of TH in 2BKT may inhibit cleavage by constraining some of the RNA in a 1-nt backtracked state that cannot be cleaved due to a mismatch between the RNA and template DNA in the pre-translocated register. However, substitutions of residues in the first helical turn of TH (β’Gln929, β’Met932, and β’Arg933) had diverse and significant effects on cleavage in 2BKT (**Figure 2,5**).

**Figure 5.**
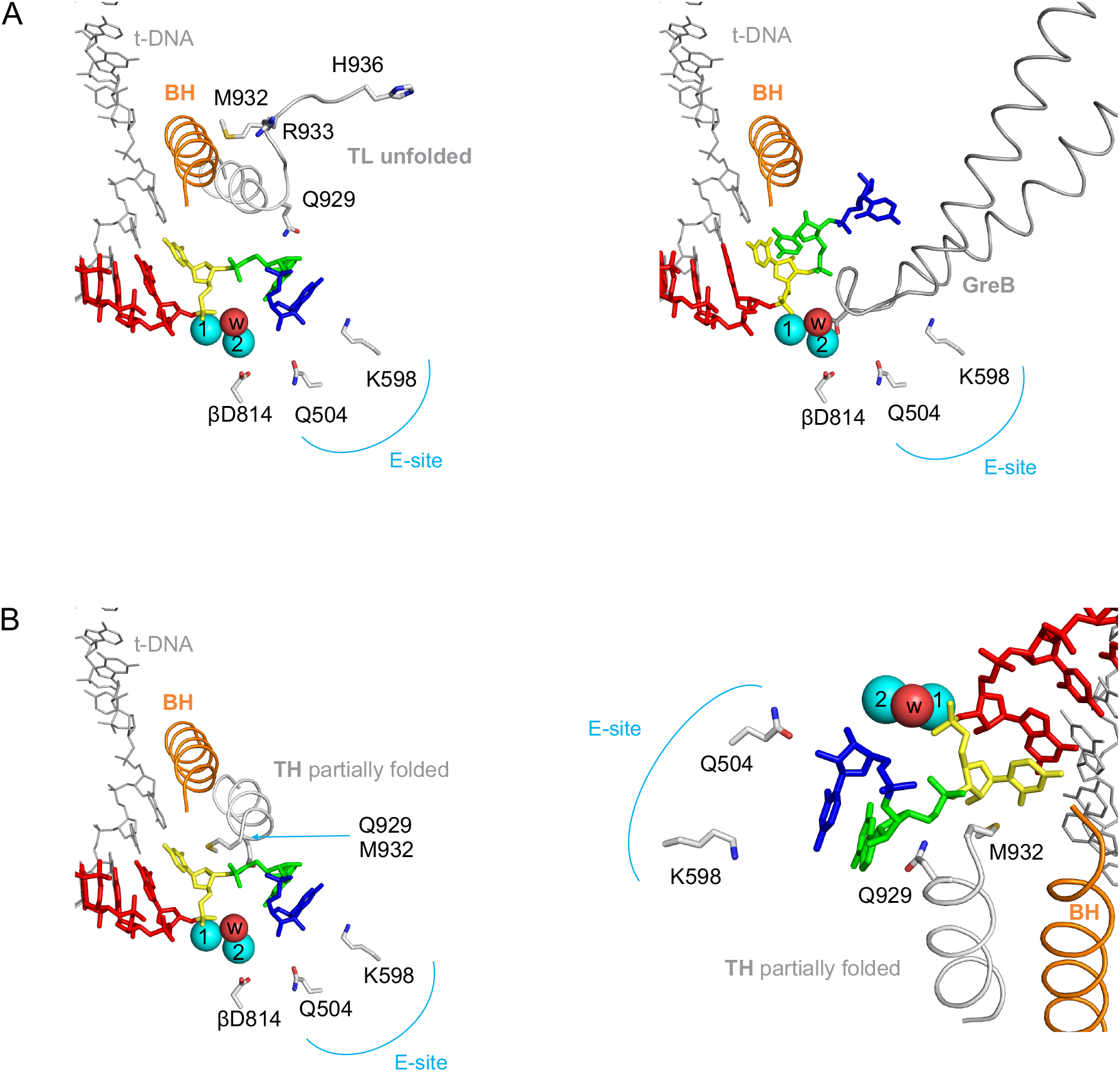
Structural models of 2-nt backtracked states of bacterial RNAP. **(A)** 2-nt backtracked state is likely incompatible with the complete folding of the TL, the backtracked RNA typically occupies the E-site (left, Zuber et al 2024). The 3’ terminal RNA nucleotide is colored blue. Binding of Gre cleavage assisting factors occupies the E-site with the protein mass and displaces the RNA towards the BH (right, Abdelkareem et al, 2019). **(B)** A limited folding of TL (up to β’Met932) is likely possible in the 2-nt backtracked state. β’Arg933 (next residue after β’Met932, not shown) is then also expected to localize close to the backtracked RNA. An overview is shown in the left panel and a closer view in the right panel.

The substitution of β’Gln929 with alanine leads to a moderate decrease in RNA cleavage rate in 2BKT, indicating that β’Gln929 likely has a modest stimulatory effect on cleavage in the 2BKT system. Replacing β’Gln929 with methionine results in a much stronger loss of function, reducing cleavage by approximately tenfold. In contrast, changing β’Gln929 to arginine leads to a significant gain of function, enhancing cleavage by more than tenfold. Similar effects were observed in both 1BKT and 2BKT. Predicting the precise mechanism by which β’Arg929 promotes RNA cleavage is challenging. The Arg side chain is long and flexible and the protein backbone at position 929 is known to twist, pointing the side chain either to the E-site (see below) or towards the main active site cavity. It is therefore possible that the effect of β’Q929R arises from either obstructing the E-site at one extremity (see below) or contacting the RNA nucleotide in the pre-translocated register at the other extremity. In the latter scenario, β’Arg929 may stimulate cleavage by altering the conformation of the sugar-phosphate backbone of the backtracked RNA including the scissile phosphate linkage.

Upon complete or partial folding of the TH, the β’Met932 residue forms stacking interactions with the nucleobase of NTP or RNA nucleotide in the pre-translocated register (Vassylyev *et al*, 2007; Basu *et al*, 2014). The substitution of β’Met932 with alanine or leucine leads to a significant loss of cleavage activity in 2BKT, suggesting that a limited folding of TH allowing such interaction may occur in 2BKT and act as a stimulus for RNA cleavage (**Figure 5B**). This hypothesis is further supported by the stimulatory effect of the β’Arg933A substitution on cleavage in 1BKT and 2BKT. β’Arg933 is the first residue in the partially folded TH that is expected to clash with the backtracked RNA (**Figure 5B**). Replacing the bulky arginine side chain with a smaller alanine could ease the partial folding of TH in 2BKT, allowing β’Met932 to stack against the nucleobase in the pre-translocated register and stimulate RNA cleavage. In contrast, the β’R933N substitution inhibits RNA cleavage, possibly because asparagine, despite being a nominally smaller side chain, can cause greater congestion in the immediate vicinity of the protein backbone than arginine.

### The role of the acidic patch in Mg^2+^ coordination

The catalysis of phosphoryl transfer reactions by RNA polymerase (RNAP) critically depends on divalent cations, particularly Mg^2+^ ions. At least two Mg^2+^ ions are essential for catalysis, and both bind near the “acidic patch” in the RNAP active site (**Figure 6A**). This patch consists of the β’ aspartate triad (Asp 460, 462, and 464) and Glu813, Asp814 contributed by the β subunit. A tightly bound Mg^2+^ ion #1 is coordinated by the aspartate triad and participates in both RNA cleavage and RNA synthesis reactions. During RNA synthesis, Mg^2+^ #1 activates the 3’OH of the RNA primer for a nucleophilic attack on the α-phosphate of the substrate NTP. A weakly bound Mg^2+^ ion #2 arrives together with the substrate NTP and is predominantly coordinated by the triphosphate moiety and Asp460 from the Asp triad. βGlu813 may also participate in coordinating Mg^2+^ #2. During the RNA cleavage reaction, Mg^2+^ #1 coordinates the substrate RNA and precisely positions it for catalysis. Mg^2+^ #2 is expected to position and activate a water molecule for nucleophilic attack on the nascent RNA. However, Mg^2+^ #2 has never been experimentally observed in the absence of substrate NTP because there are not enough ligands to coordinate it. Mg^2+^ #2 can be only approximately positioned so that a water molecule in its coordination sphere is capable of inline attack on the phosphate linkage, which would release the 3’OH-bearing RNA primer on one side and 5’ phosphorylated oligonucleotide on the other side. When positioned in such a way, Mg^2+^ #2 is located differently than during the synthesis reaction, with the most likely protein ligands being β’Asp460 and βAsp814. This hypothesis is supported by our data, as substitutions of βAsp814 were strongly inhibitory for RNA cleavage, whereas substitutions of βGlu813 were less inhibitory (**Figure 3**). The negative effects of βD814A and βD814N substitutions were particularly dramatic in 2BKT. Not only did the median cleavage time increase by 20-30-fold, but the percentage of residual uncleaved RNA also increased from 10% in wild-type RNAP to over 80% in βD814A and βD814N (**Table 2**).

**Figure 6.**
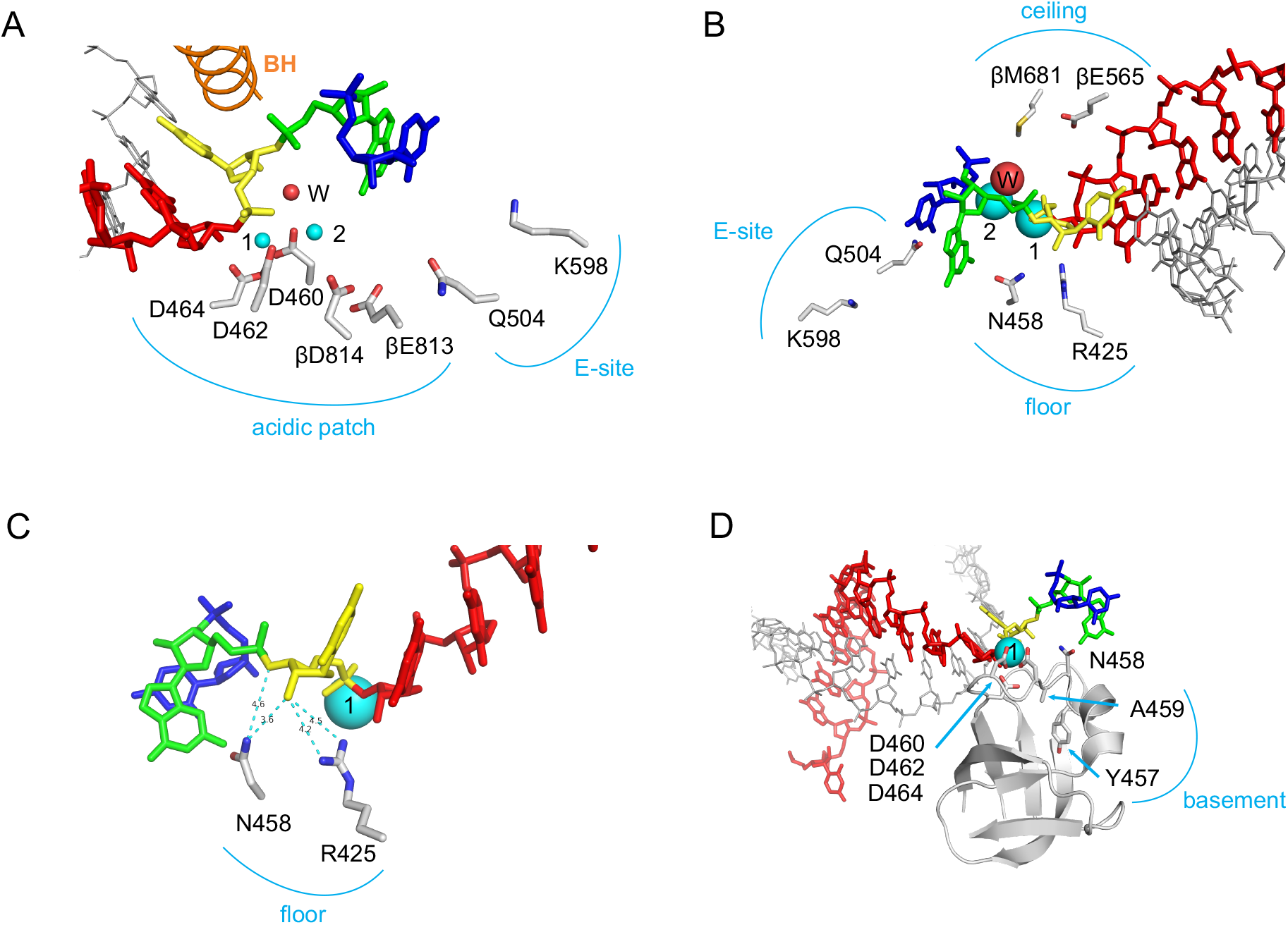
The acidic patch, ceiling, floor and the basement of the active site. **(A)** A close-up view of the acidic patch that coordinates Mg^2+^ ions. βAsp814 is well positioned to coordinate Mg^2+^ #2, whereas βGlu813 is not. **(B)** βMet681 and βGlu565 form the ceiling of the active site above the cleavage site, whereas β’Arg425 and β’Asn458 together with the β’Asp triad (460, 462, 464, not shown) form the floor of the active site below the cleavage site. **(C)** The distances between β’Arg425 and β’Asn458 and the RNA nucleotide in the pre-translocated register suggest a weak hydrogen bonding may take place in 2-nt backtracked state (PDB ID 8PID) before or after cleavage reaction. Noteworthy, these residues often approach the NTP ribose within the hydrogen bonding distance (e.g. PDB ID 4Q4Z, 2O5J). **(D)** The universally conserved β’Ala459 and semi-conserved β’Tyr457 (some species feature Phe) are part of the hydrophobic core of a beta barrel structural element that contributes the very central catalytic residues to the active site (β’Asp triad, β’Arg425 and β’Asn458). β’A459T and β’Y457F substitutions may potentially alter the conformation of β’Asn458 and/or the Asp triad.

The coordinating capacities of β’Asp460 and βAsp814 are likely insufficient to secure Mg^2+^ #2 for efficient RNA cleavage, suggesting the requirement for additional ligands. Two possibilities have been proposed. First, the RNA itself, when backtracked, could fold back into the active site and potentially chelate Mg^2+^ #2 with ribose hydroxyl groups or the nucleobase (Zenkin *et al*, 2006; Sosunova *et al*, 2013). Such scenario is more likely to take place in 1BKT than in 2BKT: a single backtracked nucleotide can fold back, but a bulkier dinucleotide is less likely to do so. Consistently, βAsp814 is more important for RNA cleavage in the latter system. Alternatively, an accessory cleavage-stimulating factor could provide the additional ligands needed to chelate Mg^2+^ #2 (Sosunova *et al*, 2003). GreB, a cleavage assisting factor, binds in the RNA polymerase active site and positions its conserved Asp residue to potentially chelate Mg^2+^ #2 in a plausible location (Abdelkareem *et al*, 2019) (**Figure 5A**). However, despite these hypothesized interactions, Mg^2+^ #2 remained invisible in the GreB bound backtracked complex.

### The effects of other residues in the main active site cavity on RNA cleavage

Our study examined the effects of four amino acid residues lining the main active site cavity of RNA polymerase in addition to the acidic patch described in the previous section. We focused on two residues lining the “ceiling” (βM681Y and βE565Q) and two residues lining the “floor” (β’R425K and β’N458S) of the active site cavity (**Figure 3, 6B**). We found that the βM681Y substitution had little effect on cleavage activity. The βE565Q substitution showed a mild stimulatory effect, but the experimental uncertainty was high. These results suggest that the ceiling of the active site has a marginal effect on cleavage activity. This conclusion is significant because, in several structures of the backtracked RNAP, backtracked RNA folds back into a region near βMet681 and βGlu565 (Wang *et al*, 2009; Sekine *et al*, 2015). Therefore, our data suggest that these structures do not represent cleavage-proficient conformations of the backtracked RNAP.

β’N458S significantly increased the cleavage rate in both 1BKT and 2BKT, while β’R425K only stimulated cleavage in 2BKT (**Figure 3**). Notably, these two residues are crucial for positioning the RNA nucleotide (or NTP) in the pre-translocated register (substrate site, A-site) (**Figure 6C**). Interestingly, most substitutions of residues that directly interact with the RNA nucleotide in the pre-translocated register (β’R425, β’N458, β’Q929, β’M932, β’R933) strongly affected the RNA cleavage rate. While the structural consequences of substitutions like β’Q929R are challenging to predict, β’R425K and β’N458S substitutions likely relax the position of the RNA nucleotide in the pre-translocated register, enabling it to adopt different conformations compared to a substrate NTP aligned for the nucleotide addition reaction (Vassylyev *et al*, 2007). These observations suggest that the optimal cleavage-proficient geometry is likely achieved when the RNA nucleotide in the pre-translocated register forms a canonical base pair with the DNA template, but its sugar-phosphate moieties are partially displaced from the canonical location characteristic for the substrate NTP or RNA 3’NMP in the pre-translocated state.

Finally, we assessed the effects of substitutions in the “basement” of the active site, β’A459T and β’Y457F on RNA cleavage activity (**Figure 3, 6D**). These substitutions are located in the hydrophobic core of the double-psi-beta barrel domain, which forms the floor of the active site and contributes several critical catalytic residues like the β’Asp triad, β’Arg425, and β’Asn458 to the active site. β’Y457F stimulated cleavage in 1BKT (∼5-fold) and had no effect in 2BKT. The substitution probably slightly alters the position of β’Asn458 that has a very large effect on cleavage in 1BKT and a smaller effect on cleavage in 2BKT. β’A459T stimulated cleavage in 2BKT (∼5-fold) and had no effect in 1BKT. The substitution probably slightly alters the arrangement of residues in the acidic patch that are responsible for coordinating Mg^2+^ ions. Mg^2+^ ions coordination becomes more important in 2BKT, hence β’A459T displays its effect only in that system.

### The role of the E-site in catalysis of the nascent RNA cleavage

The active site of the RNA polymerase has three regions: the P-site, the A-site, and the E-site (**Figure 7**). The P-site binds the 3’ NMP of the RNA primer in a post-translocated state. The A-site holds the NTP base paired to the template DNA and the 3’ NMP of the RNA primer in a pre-translocated state. The E-site is a loosely defined area in the NTP loading channel that binds NTPs in a non-sequence-specific manner (Westover *et al*, 2004; Wang *et al*, 2006; Zhang *et al*, 2014). The E-site may participate in NTP loading but is not essential. Backtracked nucleotides often localize to the E-site (Cheung & Cramer, 2011; Zuber *et al*, 2024), but RNA binding in the E-site likely inhibits cleavage activity. For instance, in 1BKT, the binding of the backtracked nucleotide in the E-site prevents it from folding back into the active site and helps chelate Mg^2+^ (see below) (Sosunova *et al*, 2013). Similarly, in 1BKT and 2BKT, the binding of backtracked nucleotides in the E-site may affect the conformation of the phosphate of the scissile phosphodiester bond in a way that inhibits cleavage. Moreover, cleavage-assisting factors are known to displace the backtracked RNA from the E-site as part of their cleavage-stimulating action (Cheung & Cramer, 2011; Abdelkareem *et al*, 2019) (**Figure 5A**). Here we report that mutations in the E-site, Q504R, and K598W (**Figures 3, 4-5**), increased the cleavage rate fivefold in 1BKT and tenfold in 2BKT. This suggests that filling the E-site with bulkier residues facilitates RNA cleavage likely by disfavoring RNA binding in the E-site.

**Figure 7.**
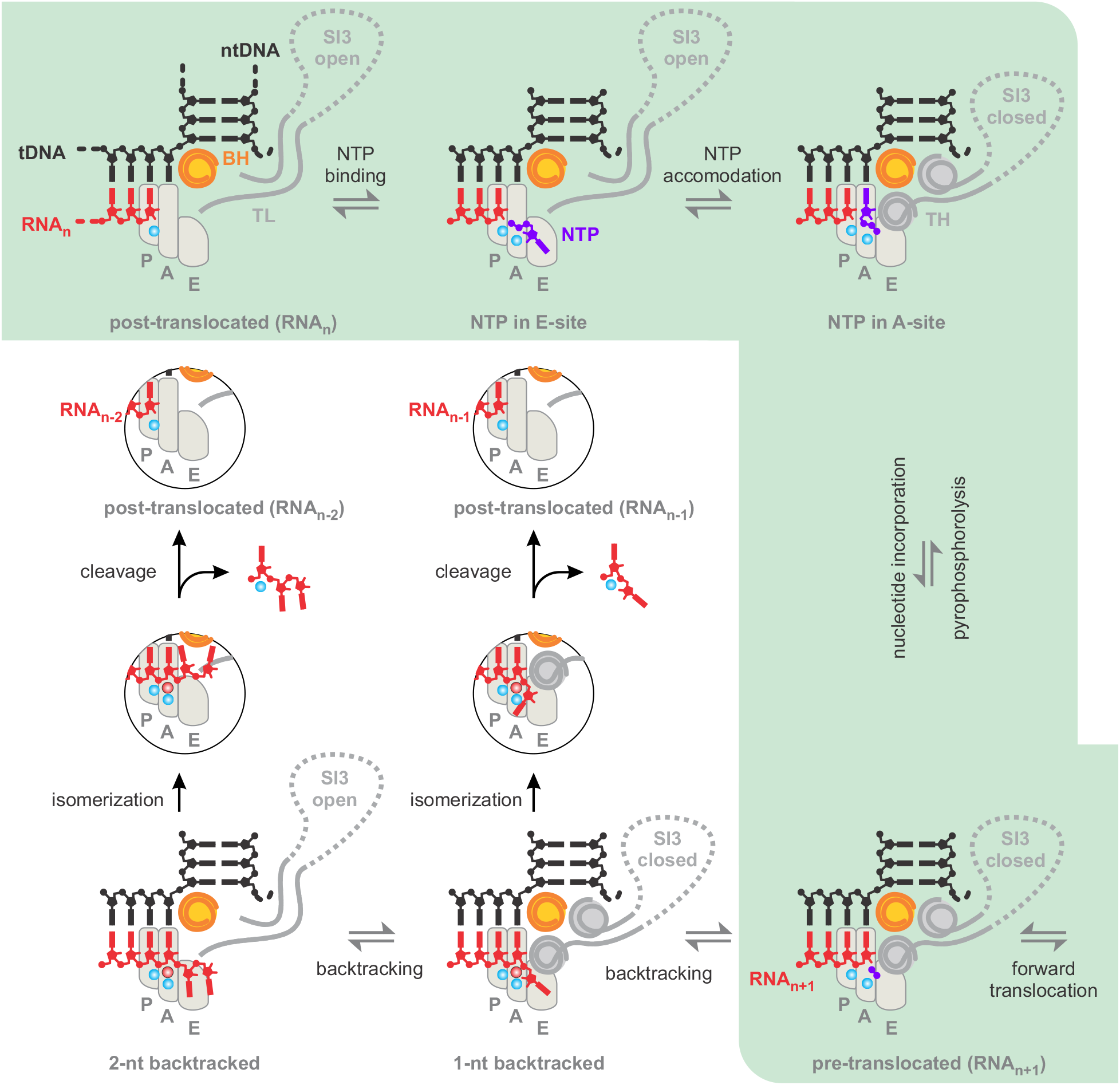
Schematics of the nucleotide addition, backtracking and RNA cleavage pathways. The areas corresponding to P-, A- and E-sites are outlined by light gray shapes. The triphosphate moiety of the substrate NTP governs binding to the E-site followed by rotation into the A-site where the complementarity to the template DNA is probed. The cognate NTP stabilizes the closed active site that positions the triphosphate moiety for efficient catalysis. Following nucleotide incorporation RNAP completes the cycle by translocating forward but may occasionally backtrack. Backtracking by 1 nt places the backtracked nucleotide into the E-site; the backtracked and the penultimate nucleotides stabilize the closed active site. Further backtracking inhibits the closure of the active site. Backtracked states can recover by forward translocation or by endonucleolytic cleavage of the nascent RNA between the A and P sites (post- and pre-translocated registers). Backtracked RNA needs to be expunged from the E-site for efficient cleavage to take place.

### General implications for the mechanism of RNA cleavage by RNAP

In general, the rate of RNA cleavage by the multisubunit RNAPs is likely affected by three major factors: (*i*) positioning the nucleophilic water molecule, (*ii*) protonating the 3’ OH group of the RNA primer, which acts as the leaving group, and (*iii*) positioning the scissile phosphate linkage to enable attack by the nucleophilic water. In intrinsic RNA cleavage, the universally conserved acidic patch likely mediates the first two effects, and the backtracked RNA may assist in some cases (Zenkin *et al*, 2006; Sosunova *et al*, 2013). In factor-assisted cleavage, accessory factors may introduce additional acidic residues, aiding in the positioning of Mg^2+^ #2, which coordinates and activates the nucleophilic water molecule (Sosunova *et al*, 2003). At the same time, the conformation of the scissile phosphate linkage can be influenced by any residue contacting the backtracked RNA in its various possible conformations, as well as by cleavage-assisting factors known to reposition the backtracked RNA (Cheung & Cramer, 2011; Abdelkareem *et al*, 2019). We suggest that most of the effects reported here are type three effects, where the RNA cleavage rate is modulated by influencing the conformation of the backtracked RNA. Substitutions in the active site cavity, TL and E-site modulate the RNA conformation by direct contacts whereas substitutions in BH and F-loop do it allosterically by influencing the TL conformation.

While conserved active site residues profoundly impact RNA cleavage, it is unreasonable to assume that these effects determined the identity of these residues. The RNA synthesis reaction, rather than RNA cleavage, played a pivotal role in shaping the identity of conserved active site residues. Consequently, the effects of conserved active site residues on the cleavage reaction are largely fortuitous. Every residue that contacts the backtracked RNA may influence its conformation and, in turn, affect RNA cleavage activity. Among the conserved active site residues, βAsp814 might be the only one that evolved specifically for RNA cleavage catalysis. Despite their fortuitous nature, the effects reported in this study provide valuable mechanistic insights. They help delineate the events surrounding the backtracked RNA during cleavage and, when combined with structural data, reveal the precise geometry of the reaction mechanism. Our results also aid in interpreting RNAP adaptations in certain species, where evolutionary pressure to regulate RNA cleavage activity may have shaped the identity of amino acid residues in specific semi-conserved positions of the TL, BH, and E-site. Lineage-specific variations at positions corresponding to the β’ subunit Ala455, Gln504, Phe773, Ile937, Ala940, and Gly1136 result in RNAPs with enhanced RNA cleavage proficiency compared to the *E. coli* enzyme (Esyunina *et al*, 2016; Riaz-Bradley *et al*, 2020). Consistently with our general inferences, lineage-specific substitutions of these residues enhance cleavage activity either by stabilizing the TH folding (F773V, I937T, A940V, and G1136Q) or by blocking the E-site (A455D, Q504R).

## Materials and methods

### Reagents and oligonucleotides

DNA and RNA oligonucleotides were purchased from Eurofins Genomics GmbH (Ebersberg, Germany) and IBA Biotech (Göttingen, Germany). DNA oligonucleotides and RNA primers are listed in **Table S1**.

### Proteins

RNAPs were expressed in *E. coli* strain T7 Express lysY/Iq (New England Biolabs, Ipswich, MA, USA) and purified by Ni-, heparin and Q-sepharose chromatography as described previously (Svetlov & Artsimovitch, 2015). RNAPs were dialyzed against a storage buffer (50% glycerol, 20 mM Tris-HCl pH 7.9, 150 mM NaCl, 0.1 mM EDTA, 0.1 mM DTT) and stored at -20 °C. Plasmids used for protein expression are listed in **Table S2**.

### TEC assembly

TECs (1 µM) were assembled by a procedure developed by Komissarova et al 2003 (Komissarova *et al*, 2003). RNA primer (1 µM) was annealed to the template DNA (1.4 µM), incubated first with RNAP (1.5 µM) for 10 min, and then with the non-template DNA (2 µM) for 20 min at 25°C. TECs were assembled in TB buffer (40 mM HEPES–KOH pH 7.5, 80 mM KCl, 5% glycerol, 0.1 mM EDTA, and 0.1 mM DTT).

### RNA cleavage reactions

Manually quenched reactions (timepoints: 15–14400 seconds) were induced by adding 2 mM MgCl2 to 100 nM TEC in TB buffer at 25 °C, and stopped at indicated times by withdrawing 20 µl samples and mixing them in 40 µl of gel loading buffer (94% formamide, 20 mM Li4-EDTA and 0.2% Orange G). RNAs were separated on 16% denaturing polyacrylamide gel and visualized with an Odyssey Infrared Imager (Li-Cor Biosciences, Lincoln, NE, USA). RNA band intensities were quantified using the ImageJ software (Abramoff *et al*, 2004).

### Data analysis

The time courses of the intrinsic RNA cleavage of the nascent RNA were fitted to the stretched exponential function. The reaction rates and median reaction times presented in **Tables 1** and **2** were calculated as in (Turtola & Belogurov, 2016). We used the stretched exponential function in the analyses because it is the simplest analytical function that robustly describes the slow processes in transcription that are, in most cases, poorly described by the single and double exponential functions (Turtola & Belogurov, 2016; Turtola *et al*, 2018).

**Table 1.** RNA cleavage rates in 1BKT complex (scaffold S270-R130-S271).

| RNAP | RNA cleavage rate $\times 10^{-3}$, s<sup>-1</sup> | | median time, min | | residual uncleaved RNA, % | |
| --- | --- | --- | --- | --- | --- | --- |
| Wild type | 1.8 | $\pm 0.6$ | 10 | $\pm 3$ | 13 | $\pm 1$ |
| $\beta'$P750L | 8.3 | $\pm 0.8$ | 2.0 | $\pm 0.2$ | 17 | $\pm 3$ |
| $\beta'$F773V | 10 | $\pm 5$ | 1.9 | $\pm 0.9$ | 16 | $\pm 2$ |
| $\beta'$Q929A | 1.1 | $\pm 0.1$ | 15 | $\pm 1$ | 29 | $\pm 1$ |
| $\beta'$Q929M | 0.4 | $\pm 0.1$ | 40 | $\pm 9$ | 37 | $\pm 0.02$ |
| $\beta'$Q929R | 25 | $\pm 5$ | 0.66 | $\pm 0.12$ | 9 | $\pm 5$ |
| $\beta'$M932A | 0.9 | $\pm 0.2$ | 19 | $\pm 5$ | 19 | $\pm 0.4$ |
| $\beta'$M932L | 1.2 | $\pm 0.2$ | 14 | $\pm 2$ | 18 | $\pm 3$ |
| $\beta'$R933A | 11 | $\pm 0.2$ | 1.46 | $\pm 0.03$ | 15 | $\pm 5$ |
| $\beta'$R933N | 0.6 | $\pm 0.3$ | 34 | $\pm 20$ | 21 | $\pm 3$ |
| $\beta'$H936A | 2.7 | $\pm 0.3$ | 6.2 | $\pm 0.6$ | 22 | $\pm 2$ |
| $\beta'$H936Q | 1.8 | $\pm 0.9$ | 10 | $\pm 5$ | 23 | $\pm 1$ |
| $\beta'$I937T | 4.3 | $\pm 0.5$ | 3.9 | $\pm 0.5$ | 19 | $\pm 8$ |
| $\beta'$T1135V | 1.6 | $\pm 0.4$ | 10 | $\pm 2$ | 24 | $\pm 6$ |
| $\beta'$G1136S | 8 | $\pm 2$ | 2.2 | $\pm 0.6$ | 13 | $\pm 3$ |
| $\beta'$ ΔSI3 | 7 | $\pm 3$ | 2.7 | $\pm 1.2$ | 20 | $\pm 7$ |
| βE565Q | 6 | $\pm 2$ | 3.1 | $\pm 1.1$ | 8.4 | $\pm 2.4$ |
| βM681Y | 2.9 | $\pm 0.3$ | 5.9 | $\pm 0.6$ | 27 | $\pm 0.4$ |
| βE813A | 1.1 | $\pm 0.6$ | 17 | $\pm 9$ | 12 | $\pm 1$ |
| βE813Q | 0.61 | $\pm 0.01$ | 27.3 | $\pm 0.3$ | 39 | $\pm 5$ |
| βD814A | 0.38 | $\pm 0.08$ | 45 | $\pm 9$ | 25 | $\pm 5$ |
| βD814N | 0.17 | $\pm 0.07$ | 108 | $\pm 44$ | 40 | $\pm 10$ |
| $\beta'$R425K | 2.9 | $\pm 0.8$ | 5.9 | $\pm 1.7$ | 8.1 | $\pm 1.9$ |
| $\beta'$R425L | 0.43 | $\pm 0.13$ | 41 | $\pm 13$ | 18 | $\pm 10$ |
| $\beta'$Y457F | 4.9 | $\pm 0.6$ | 3.4 | $\pm 0.4$ | 17 | $\pm 5$ |
| $\beta'$N458S | 500 | $\pm 260$ | 0.04 | $\pm 0.02$ | 5.5 | $\pm 2.2$ |
| $\beta'$A459T | 1.1 | $\pm 0.4$ | 16 | $\pm 6$ | 13 | $\pm 4$ |
| $\beta'$Q504R | 11 | $\pm 5$ | 1.7 | $\pm 0.8$ | 14 | $\pm 3$ |
| $\beta'$K598W | 14 | $\pm 5$ | 1.2 | $\pm 0.4$ | 14 | $\pm 3$ |

**Table 2.** RNA cleavage rates in 2BKT complex (scaffold S290-R124-S291).

| RNAP | RNA cleavage rate $\times 10^{-3}$, s<sup>-1</sup> | | median time, min | | residual uncleaved RNA, % | |
| --- | --- | --- | --- | --- | --- | --- |
| Wild type | 1.3 | $\pm 0.2$ | 12.8 | $\pm 2.4$ | 12 | $\pm 3$ |
| $\beta$'P750L | 0.66 | $\pm 0.03$ | 25 | $\pm 1$ | 35 | $\pm 21$ |
| $\beta$'F773V | 0.47 | $\pm 0.15$ | 38 | $\pm 12$ | 18 | $\pm 6$ |
| $\beta$'Q929A | 0.47 | $\pm 0.04$ | 35 | $\pm 3$ | 40 | $\pm 2$ |
| $\beta$'Q929M | 0.13 | $\pm 0.03$ | 138 | $\pm 38$ | 67 | $\pm 2$ |
| $\beta$'Q929R | 19 | $\pm 1$ | 0.86 | $\pm 0.04$ | 8.7 | $\pm 0.5$ |
| $\beta$'M932A | 0.24 | $\pm 0.02$ | 69 | $\pm 6$ | 27 | $\pm 6$ |
| $\beta$'M932L | 0.27 | $\pm 0.07$ | 63 | $\pm 16$ | 28 | $\pm 6$ |
| $\beta$'R933A | 6.3 | $\pm 1.4$ | 2.7 | $\pm 0.6$ | 18 | $\pm 5$ |
| $\beta$'R933N | 0.23 | $\pm 0.01$ | 73 | $\pm 3$ | 28 | $\pm 8$ |
| $\beta$'H936A | 1.7 | $\pm 0.2$ | 10 | $\pm 1$ | 26 | $\pm 1$ |
| $\beta$'H936Q | 1.09 | $\pm 0.01$ | 15.3 | $\pm 0.2$ | 37 | $\pm 15$ |
| $\beta$'I937T | 1.26 | $\pm 0.03$ | 13.2 | $\pm 0.4$ | 23 | $\pm 6$ |
| $\beta$'T1135V | 0.26 | $\pm 0.01$ | 64 | $\pm 1$ | 41 | $\pm 6$ |
| $\beta$'G1136S | 0.54 | $\pm 0.05$ | 31 | $\pm 3$ | 27 | $\pm 12$ |
| $\beta$'ΔSI3 | 0.64 | $\pm 0.01$ | 26 | $\pm 0.2$ | 23 | $\pm 9$ |
| βE565Q | 6.9 | $\pm 4.2$ | 2.9 | $\pm 1.8$ | 3.8 | $\pm 0.2$ |
| βM681Y | 0.64 | $\pm 0.15$ | 27 | $\pm 6$ | 29 | $\pm 12$ |
| βE813A | 0.26 | $\pm 0.21$ | 98 | $\pm 80$ | 27 | $\pm 16$ |
| βE813Q | 0.19 | $\pm 0.04$ | 91 | $\pm 19$ | 51 | $\pm 14$ |
| βD814A | 0.04 | $\pm 0.02$ | 452 | $\pm 205$ | 85 | $\pm 2$ |
| βD814N | 0.06 | $\pm 0.01$ | 282 | $\pm 48$ | 81 | $\pm 11$ |
| $\beta$'R425K | 13 | $\pm 0.2$ | 1.27 | $\pm 0.02$ | 16 | $\pm 1$ |
| $\beta$'R425L | 0.92 | $\pm 0.11$ | 18.3 | $\pm 2.2$ | 16 | $\pm 8$ |
| $\beta$'Y457F | 1.3 | $\pm 0.3$ | 12.6 | $\pm 2.4$ | 21 | $\pm 3$ |
| $\beta$'N458S | 11 | $\pm 1$ | 1.5 | $\pm 0.1$ | 10 | $\pm 9$ |
| $\beta$'A459T | 7.6 | $\pm 3$ | 2.4 | $\pm 1.1$ | 10 | $\pm 2$ |
| $\beta$'Q504R | 13 | $\pm 2$ | 1.3 | $\pm 0.2$ | 20 | $\pm 3$ |
| $\beta$'K598W | 24.1 | $\pm 0.1$ | 0.69 | $\pm 0.01$ | 24 | $\pm 7$ |

## Data availability

All data are included in the article and supporting information file.

## Supporting information

This article contains supporting information.

## Acknowledgements

The authors would like to thank Dr. Irina Artsimovitch for providing RNA polymerase expression plasmids for variants β’N458S, βE813A and β’R933A. Authors thank Dr. Anssi Malinen, Dr. Matti Turtola, Dr. Thadee Grocholski, Dr. Ranjit Prajapati, Eeva Vieras and Karina Šapovalovaitė for constructing expression plasmids.

## Declaration of Generative AI and AI-assisted technologies in the writing process

During the preparation of this work the authors used Gemini to improve the language and readability. After using these tools, the authors reviewed and edited the content as needed and take full responsibility for the content of the publication.

## Supplementary information

**Table S1.**
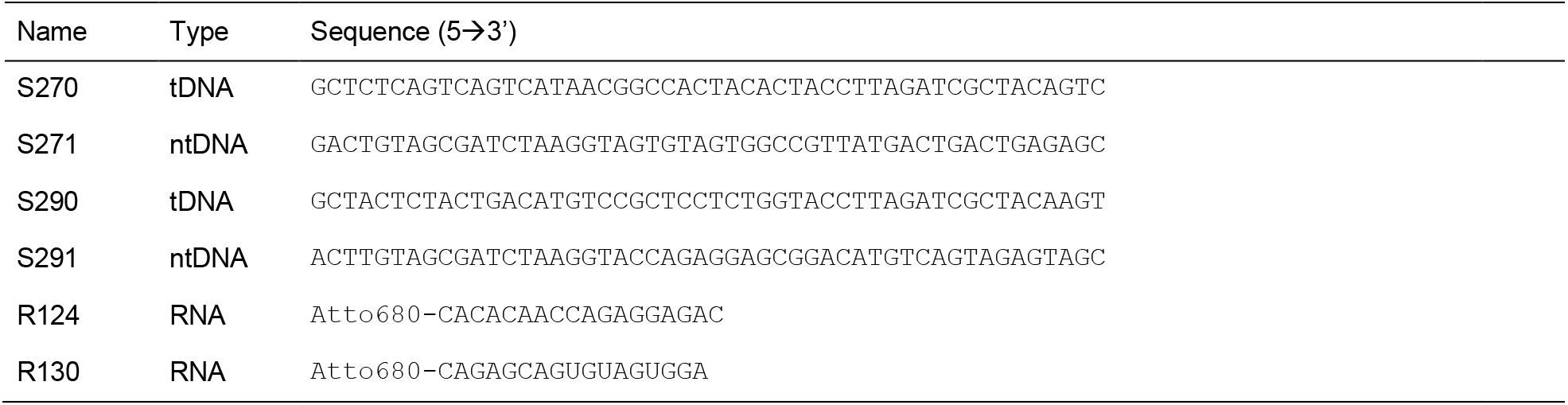
DNA and RNA oligonucleotides.

**Table S2.** RNAP expression vectors.

| Name | Description | Source/reference |
| --- | --- | --- |
| pVS10 | *E. coli* wild-type RNAP (T7p-α-β-$\beta'$-His<sub>6</sub>-ω) | Belogurov *et al*, 2007 |
| pAM012 | $\beta'$M932A RNAP (T7p-α-β-$\beta'$[M932A]-TEV_His<sub>10</sub>-ω) | Malinen *et al*, 2012 |
| pAM013 | $\beta'$Q929A RNAP (T7p-α-β-$\beta'$[Q929A]-TEV_His<sub>10</sub>-ω) | Turtola *et al*, 2018 |
| pAM014 | $\beta'$M932L RNAP (T7p-α-β-$\beta'$[M932L]-TEV_His<sub>10</sub>-ω) | Turtola *et al*, 2018 |
| pAM017 | $\beta'$R425L RNAP (T7p-α-β-$\beta'$[R425L]-TEV_His<sub>10</sub>-ω) | this work |
| pAM018 | $\beta'$R425K RNAP (T7p-α-β-$\beta'$[R425K]-TEV_His<sub>10</sub>-ω) | Mäkinen *et al*, 2021 |
| pEV001 | βE813Q RNAP (T7p-α-His<sub>6</sub>-β[E813A]-$\beta'$-ω) | this work |
| pEV002 | βD814N RNAP (T7p-α-His<sub>6</sub>-β[E813A]-$\beta'$-ω) | this work |
| pGB130 | $\beta'$H936A RNAP (T7p-α-β-$\beta'$[H936A]-TEV_His<sub>10</sub>-ω) | Malinen *et al*, 2012 |
| pIA528+pIA839 | $\beta'$N458S RNAP (T7p-α-β-$\beta'$[N458S]-His<sub>6</sub>) + *araBp*-ω | Svetlov *et al*, 2004 |
| pIA624+pIA839 | βE813A RNAP (T7p-α-His<sub>6</sub>-β[E813A]-$\beta'$) + *araBp*-ω | Vassilyev *et al*, 2005 |
| pIA846 | $\beta'$R933A RNAP (T7p-α-β-$\beta'$[R933A]-His<sub>6</sub>-ω) | Artsimovitch *et al*, 2011 |
| pJM001 | $\beta'$G1136S RNAP (T7p-α-β-$\beta'$[G1136S]-TEV_His<sub>10</sub>-ω) | Turtola *et al*, 2018 |
| pJM002 | $\beta'$H936Q RNAP (T7p-α-β-$\beta'$[H936Q]-TEV_His<sub>10</sub>-ω) | Turtola *et al*, 2018 |
| pJM009 | $\beta'$T1135V RNAP (T7p-α-β-$\beta'$[T1135V]-TEV_His<sub>10</sub>-ω) | this work |
| pJM011 | βE565Q RNAP (T7p-α-His<sub>6</sub>-β[E565Q]-$\beta'$-ω) | this work |
| pJM014 | $\beta'$Q504R RNAP (T7p-α-β-$\beta'$[Q504R]-TEV_His<sub>10</sub>-ω) | Turtola *et al*, 2018 |
| pJM016 | $\beta'$K598W RNAP (T7p-α-β-$\beta'$[K598W]-TEV_His<sub>10</sub>-ω) | Turtola *et al*, 2018 |
| pJM017 | $\beta'$Q929M RNAP (T7p-α-β-$\beta'$[Q929M]-TEV_His<sub>10</sub>-ω) | Mäkinen *et al*, 2021 |
| pJM018 | $\beta'$I937T RNAP (T7p-α-β-$\beta'$[I937T]-TEV_His<sub>10</sub>-ω) | this work |
| pKS001 | βD814A RNAP (T7p-α-His<sub>6</sub>-β[D814A]-$\beta'$-ω) | this work |
| pKS003 | $\beta'$Y457F RNAP (T7p-α-β-$\beta'$[Y457F]-TEV_His<sub>10</sub>-ω) | this work |
| pMT024 | βM681Y RNAP (T7p-α-His<sub>6</sub>-β[M681Y]-$\beta'$-ω) | Turtola *et al*, 2018 |
| pMV013 | $\beta'$A459T RNAP (T7p-α-β-$\beta'$[A459T]-TEV_His<sub>10</sub>-ω) | this work |
| pMV018 | $\beta'$Q929R RNAP (T7p-α-β-$\beta'$[Q929R]-TEV_His<sub>10</sub>-ω) | this work |
| pRP010 | $\beta'$R933N RNAP (T7p-α-β-$\beta'$[R933N]-TEV_His<sub>10</sub>-ω) | this work |
| pVS10-ΔSI3(b) | $\beta'$ ΔSI3(b) RNAP (T7p-α-β-$\beta'$[Δ945-1132]-His<sub>6</sub>-ω) | Esyunina *et al*, 2016 |
| pTG012 | $\beta'$P750L RNAP (T7p-α-β-$\beta'$[P750L]-TEV_His<sub>10</sub>-ω) | Malinen *et al*, 2014 |
| pVS48 | $\beta'$F773V RNAP (T7p-α-β-$\beta'$[F773V]-His<sub>6</sub>-ω) | Svetlov *et al*, 2007 |

